# Grouping effects in numerosity perception under prolonged viewing conditions

**DOI:** 10.1101/460741

**Authors:** Leo Poom, Marcus Lindskog, Anders Winman, Ronald van den Berg

## Abstract

Humans can estimate numerosities – such as the number sheep in a flock – without deliberate counting. A number of biases have been identified in these estimates, which seem primarily rooted in the spatial organization of objects (grouping, symmetry, etc). Most previous studies on the number sense used static stimuli with extremely brief exposure times. However, outside the laboratory, visual scenes are often dynamic and freely viewed for prolonged durations (e.g., a flock of moving sheep). The purpose of the present study is to examine grouping-induced numerosity biases in stimuli that more closely mimic these conditions. To this end, we designed two experiments with limited-dot-lifetime displays (LDDs), in which each dot is visible for a brief period of time and replaced by a new dot elsewhere after its disappearance. The dynamic nature of LDDs prevents subjects from counting even when they are free-viewing a stimulus under prolonged presentation. Subjects estimated the number of dots in arrays that were presented either as a single group or were segregated into two groups by spatial clustering, dot size, dot color, or dot motion. Grouping by color and motion reduced perceived numerosity compared to viewing them as a single group. Moreover, the grouping effect sizes between these two features were correlated, which suggests that the effects may share a common, feature-invariant mechanism. Finally, we find that dot size and total stimulus area directly affect perceived numerosity, which makes it difficult to draw reliable conclusions about grouping effects induced by spatial clustering and dot size. Our results provide new insights into biases in numerosity estimation and they demonstrate that the use of LDDs is an effective method to study the human number sense under prolonged viewing.

## INTRODUCTION

Humans can estimate the quantity of a set of objects without explicitly counting them. Lately, this cognitive ability has received overwhelming attention in psychological research, much due to a suggested link between the acuity of human number sense and performance on arithmetic tasks [1], as well as the proposal of a dedicated approximate number system to explain this link [2]. However, research on the number sense dates back to as early as the 1870s, when W. Stanley Jevons found that the error in numerosity estimates increases with the number of beans landing in the box. This finding has been replicated extensively and is now known to be consistent with Weber’s law [3]. In the 150 years of research following the study by Jevons, a wide range of biases have been identified in the number sense, many of which are related to the spatial arrangement of objects (see Fig. 1 for an example). However, people seem to have known of the existence of interactions between number sense and spatial arrangement long before they became subject of scientific investigation. A famous example, noted by Ginsburg [4], is a story in the Genesis contains in which it is told that Jacob divided the animals that he gifted to his brother Esau into smaller groups, with the intention to make the gift appear more numerous. Several studies have found evidence in favor of Jacob’s hypothesis that perceived numerosity increases with the number of groups [5,6], but at least one other study has found an opposite effect [7]. Another class of studies has examined effects of spatial regularity and found that, in general, regularly spaced items are perceived as more numerous than randomly arranged, more clustered items [4,8–11]. Furthermore, both spatial and temporal order effects on perceived numerosity have been reported: stimuli presented on the left are on average perceived as more numerous than stimuli on the right [12] and when two stimuli are presented in sequence, the former tends to be perceived as less numerous than the latter [13]. In addition, it has been found that the total stimulus area and – conversely – object density affects numerosity estimates [14,15]: the more spread out a set of objects, the more numerous they generally appear to be. Finally, it has been found that reducing the symmetry in a stimulus increases both perceived numerosity [16] and numerosity estimation accuracy [17], while physically connecting objects with line segments decreases perceived numerosity [18,19]. Besides these effects of spatial arrangement, several studies have found that perceived numerosity also depends on the size of the individual objects, although with mixed results: some of these studies report that larger elements are perceived as more numerous than smaller ones [20–23], while others report the opposite effect [4,24–28].

**Figure 1.**
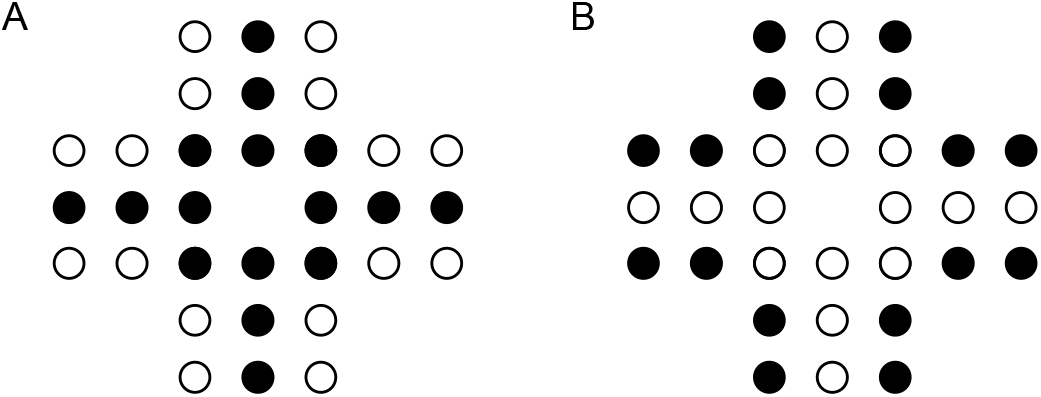
The solitaire illusion. (A) While the figure contains equal amounts of black and white dots, the black ones appear to be more numerous to most observers. (B) Reversing the colors results in an opposite effect, which indicates that the relative overestimation of the central items is due to the spatial arrangement of the objects and not due to their color.

### Numerosity judgments in the laboratory versus numerosity judgments in the wild

Many of the methodological aspects have hardly changed since Jevons’s landmark study [29]. In particular, most experiments still use static stimuli with extremely brief stimulus exposures. While this approach has revealed valuable insights into basic aspects of the human number sense, it is unclear how representative they are for situations outside the laboratory. For example, when estimating the number of people in a crowd or the number of birds in a flock, the relevant visual information is usually dynamic and available for an extended duration. Indeed, outside the laboratory, scene dynamics – and not exposure time – often seems to be the key limiting factor when estimating numerosities. Since prolonged presentation might be necessary to accomplish reliable perceptual grouping [30,31] and possibly reduces bottom-up influences of basic stimulus features, it is unclear whether biases and other findings found using brief displays generalize directly to situations with prolonged stimulus viewing.

### Study aims

The aim of the present study is to explore perceptual grouping effects on perceived numerosity in dynamic stimuli that are freely viewed for prolonged durations. To this end, we develop an experimental paradigm with limited-dot-liftetime displays (LDDs), in which a specified number of dots are visible at any moment, but each individual dot lives for only a brief duration. When a dot disappears, a new dot appears at another location. The dynamic nature of LDDs makes it impossible to count dots outside the subitizing range, even under prolonged viewing of the stimulus. Moreover, the dynamics make that the perceived stimulus area is close to the area specified by the experimenter, which removes unwanted trial-to-trial variability in perceived area, which is typically large when using static stimuli. We use this paradigm to measure perceived numerosity in arrays in which objects either form a single group or are segregated into two groups.

## EXPERIMENT 1

### Methods

#### Availability of data and analysis files

The data and JASP analysis files related to this experiment are available at https://osf.io/vbdf5/.

#### Participants

Fifty-six adults were recruited from the student population at the department of Psychology at Uppsala University (23±2.3 years of age, 31 females). Participants received either course credit or a cinema ticket for their participation. One participant was excluded from the analyses, because of partial loss of data due to a technical error.

#### Stimuli and procedure

Each test array consisted of a set of dots and was presented on the left side of the screen. A response array with an adjustable number of dots was simultaneously presented on the right side of the screen (Fig. 2A). Participants were instructed to adjust the number of dots in the response array to match the numerosity of the test array. The arrays were horizontally separated by 8 degrees of visual angle and viewed at a distance of approximately 60 cm. A vertical dashed line was shown at the center of the screen to separate the screen into two stimulus areas. Dots in both arrays were presented within areas of 8 × 8 visual degrees, except when grouping was based on spatial clustering (see below). Dots were randomly scattered within the stimulus area, with the constraint that the minimum center-to-center distance between each pair of dots was 0.80 deg. All dots had a lifetime of 300 milliseconds, with randomized temporal phases to avoid simultaneous replacement. Simultaneously with the disappearance of a dot, another one would appear elsewhere.

**Figure 2.**
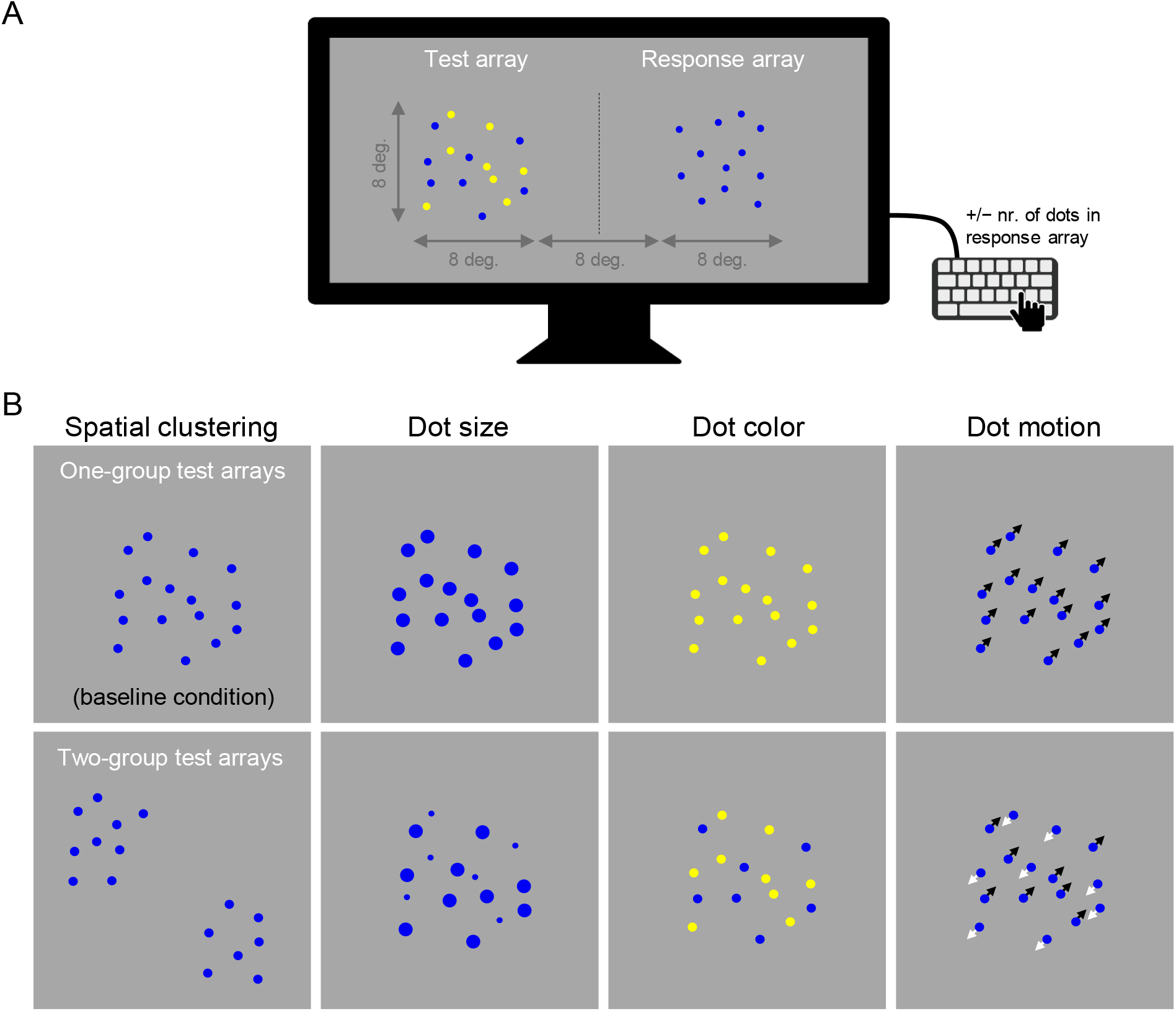
Stimulus examples in Experiment 1. (A) A snapshot of an example stimulus in Experiment 1 (not to scale). In this example, the test array is composed of two groups, with dot color being the grouping feature. Subjects adjusted the number of dots in the response array through key presses. (B) Snapshots of the eight types of test array used in Experiment 1 (example videos of these stimuli are provided at https://osf.io/vbdf5/). In the one-group conditions (top row), test dots appeared as a single group. In the two-group conditions (bottom row), the dot array was segregated into two groups, based on spatial clustering, dot size, dot color, or dot motion. The visual properties of the adjustable response array were the same throughout the experiment and matched the properties of dots in the one-group spatial clustering condition (top left). Therefore, we refer to this condition as a baseline condition.

The experiment contained four one-group conditions (Fig. 2B, top row). In the first of these, the visual properties of the test array were identical to those of the response array (Fig. 2B, top, left panel). We occasionally refer to this condition as the *baseline* condition. In the other three one-group conditions, dots in the test array differed from dots in the response array by their size (1.0 deg. *vs*. 0.50 deg. in the response array), color (yellow *vs*. blue in the response array), or motion (1.5 deg. per second *vs*. stationary in the response array).

In addition, the experiment contained four two-group conditions, in which the dots in the test array were divided into groups by spatial clustering, dot size, dot color, or dot motion (Fig. 2B, bottom). These grouping features are expected to cause strong perceptual grouping effects, as established by the Gestalt laws [32] of proximity, common size, common color, and common motion. In the two-group *spatial clustering condition*, dots were presented within two separate square areas of 32 deg^2^ each, presented with a center-to-center separation of 8 deg. To avoid location-specific effects, the entire test stimulus was rotated with a random angle around the midpoint between the two groups. In the two-group *size condition*, dots in one group were smaller (Ø=0.25 deg.) and dots in the other group larger (Ø=1.0 deg.) than dots in the response array (Ø=0.50 deg.). In the two-group *color condition*, dots in one group were yellow and dots in the other group blue. Finally, in the two-group *motion condition*, dots in one group travelled with a speed of 1.5 deg. per second in a random direction and dots in the other group travelled with the same speed in the opposite direction. On each two-group trial, one of the two groups was assigned half of the total number of dots plus or minus 0, 1, or 2, dots (randomly chosen) and the remaining dots were assigned to the other group. In each of the 8 described conditions, the test stimulus contained 16 dots on half of the trials and 20 on the other half.

The visual properties of the response array were fixed throughout the experiment: the dots were blue, 0.50 deg. in diameter, stationary during their lifetime, and presented as a single group. The initial number of dots in the response array was on each trial drawn from a uniform distribution on integers 8-24 when the test stimulus contained 16 dots and on integers 10-30 when the test stimulus contained 20 dots. Adjustments in the response stimulus were made by pressing the “F” and “K” keys to respectively decrease or increase the number of dots. The number of dots in the response array was constrained to the range 0 to 50^a^. Responses were submitted by pressing space bar. No feedback was provided. Stimulus arrays were presented in random order with 8 repetitions, giving a total of 128 trials (8 conditions × 2 numerosities × 8 repetitions). Videos of stimulus examples are available at https://osf.io/vbdf5/.

#### Analyses

The dependent variable in all analyses is the final number of dots in the response array (averaged across trial repetitions), which we refer to as the point of subjective equality (PSE). We analyze the data using ANOVAs, t-tests, and Pearson correlation tests. In addition to frequentist *p* values we also report various types of Bayes factors. The first type, denoted BF_10_, specifies the ratio between the evidence for a hypothesis H_1_ relative to another hypothesis H_0_, where the latter typically is the “null” hypothesis of no effect. E.g., a finding of BF_10_=3 means that the data are 3 times more likely under the hypothesis that there is an effect (H_1_) compared to the hypothesis that the effect size is 0 (H_0_). When doing a directed test, we denote the Bayes factor as BF_+0_ (when “H_1_: Group 1 > Group 2”) or BF_−0_ (when “H_1_: Group 1 < Group 2”). Finally, in the case of an ANOVA, BF_inclusion_ denotes the evidence for an effect averaged across all hypotheses that include the effect relative to all hypotheses that do not include the effect^b^. For example, BF_inclusion_=3 for a main effect of some factor *F*_1_ in a multi-factor ANOVA indicates that the data are on average 3 times more likely under hypotheses that include a main effect of *F*_1_ than under hypotheses without this main effect. We use the convention provided by Wagenmakers et al. [33] to label the strength of evidence provided by a Bayes factor as “extreme” (BF>100), “very strong” (30<BF≤100), “strong” (10<BF≤30), “moderate” (3<BF≤10), “anecdotal” (1<BF≤3), or “none” (BF=1). All analyses were performed using the JASP software package [34].

### Results

Subjects matched the number of dots in an adjustable response array to the number of dots in a test array. Dots in the test array were either presented as a single group or segregated into two groups by spatial clustering, dot size, dot color, or dot motion (Fig. 2B). The aim of the experiment is to examine feature-driven effects and grouping-driven effects on perceived numerosity. With “feature-driven effects”, we refer to differences in perceived numerosity that occur due to a feature difference between two conditions. These effects will be examined by comparing the one-group baseline condition – in which the test stimulus had the same visual properties as the response array (Fig. 2B, top left) – with the one-group conditions in which the test stimulus differed from the response array in dot size, dot color, or dot motion. With “grouping-driven effects”, we refer to differences in perceived numerosity that occur due to a difference in the number of perceptual groups between two conditions. These effects will be examined by comparing two-group conditions with corresponding one-group conditions. However, a complication in the latter analysis is that grouping-driven effects can only be reliably assessed when no feature-driven effect was found. The reason for this is that two-group arrays necessarily differ both in terms of its basic visual features and in the number of groups from one-group arrays (for example, when using dot size to segregate dots into two groups, the two-group arrays contain dots with two different sizes while one-group arrays contain dots with a single size). Therefore, if a feature-driven effect is found, then any difference in perceived numerosity between two-group arrays and one-group arrays may be a purely feature-driven effect or a combination of feature-driven and grouping-driven effects. We present the results per feature, starting with the features in which we found no feature-driven effects.

#### Motion

The average reported number of dots is nearly identical between the one-group motion condition (Fig. 3, right graph, filled circles) and the stationary one-group baseline condition (Fig. 3, right graph, open circles). Indeed, t-tests provide moderately strong evidence in favor of the null hypothesis, both in trials with 16 dots (Δ=0.14, BF_10_=0.18, *p*=0.26) and in trials with 20 dots (Δ=0.19, BF_10_=0.19, *p*=0.24). This indicates that the presence of motion by itself did not affect perceived numerosity and, therefore, that any difference that we may find between one-group and two-group conditions is likely to be due to grouping. We find that the average number of reported dots in the two-group motion condition is lower than in the one-group motion condition, both in trials with 16 dots (Δ=−0.22) and in trials with 20 dots (Δ=−0.86). A statistical analysis reveals extremely strong evidence for a decrease in trials with 20 dots (BF_−0_=2.06·10^2^, *p*<.001), but results for trials with 16 dots are inconclusive (BF_−0_=0.43, *p*=.15). In summary, motion by itself does not seem to affect perceived numerosity, but when using motion to segregate dots into groups, there is evidence that the average perceived number of dots decreases.

**Figure 3.**
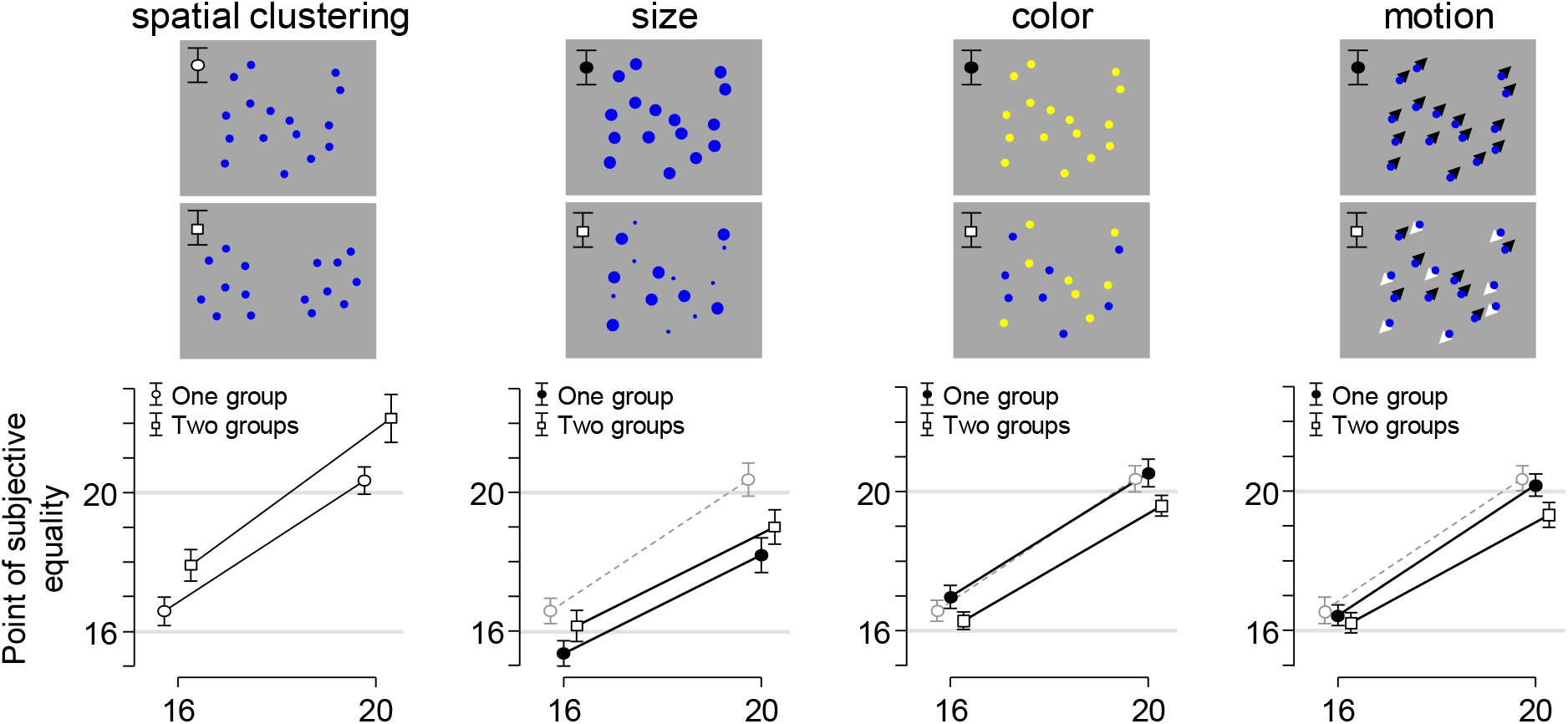
Subject-averaged numerosity estimates across the 8 stimulus conditions in Experiment 1. Subjects matched the number of dots in an adjustable response array to the number of dots in 8 different types of test array. We refer to the estimates as the point of subjective equality. The visual properties of test array in the one-group spatial clustering condition (top left) were identical to those of the response array. For comparison, we replotted the PSEs in this condition (black circles in left graph) in the results for the size, color, and motion conditions (gray circles). Segregating dots into two groups resulted in an increase in perceived numerosity in the spatial clustering and dot-size conditions and in a decrease in the color and motion conditions. Error bars indicate 95% confidence intervals.

#### Color

The results for color are very similar to the results for motion. The average reported number of dots in one-group trials with yellow arrays was nearly identical to the average number of reported dots in the baseline condition (Fig. 3, third graph from the left; compare filled circles with open circles). Although the results for trials with 16 dots are inconclusive (Δ=0.41, BF_10_=0.67, *p*=.96), we find moderate evidence for the null hypothesis in trials with 20 dots (Δ=0.16, BF_10_=0.17, *p*=.71). Hence, we can be reasonably confident that any difference that we may find between two-group trials and one-group trials is likely to be grouping-driven rather than feature-driven. Just as was the case with motion-based grouping, we find that numerosity estimates were on average lower in trials where dots were grouped by color (Fig. 3, third column, squares) compared to one-group trials (Fig. 3, third column, filled circles). A t-test provides strong evidence for this effect in trials with 16 dots (Δ=−0.69, BF_−0_=66.6, *p*<.001) and extremely strong evidence in trials with 20 dots (Δ=−0.94, BF_−0_=45.5·10^2^, *p*<.001). In summary, the results for color are similar to the results for motion: this feature does not directly affect perceived numerosity, but when it is used to segregate a set of dots into two groups, the average reported number of dots decreases.

#### Dot size

To examine whether dot size directly affected perceived numerosity, we compare estimates in the one-group baseline condition (Fig. 2B, top left) with estimates in the one-group condition with larger dots (Fig. 2B, top, second column). This comparison reveals extremely strong evidence for that perceived numerosity was smaller in the condition with larger dots, both in trials with 16 dots (Δ=−1.22, BF_−0_=7.89·10^3^,*p*<.001) and in trials with 20 dots (Δ=−2.18, BF_−0_=3.66·10^5^,*p*<.001). As explained above, this feature-driven effect makes it difficult to assess grouping-driven effects, because any effect that we may find between two-group trials and one-group trials could be either due to dot size differences or due to a combination of dot size differences and a difference in the number of groups.

The number of reported dots in the two-group trials (Fig. 3, squares in second graph) is on average larger than in one-group trials with only large dots (Fig. 3, filled circles in second graph). A t-test reveals moderately strong statistical evidence for a difference on both the trials with 16 dots (Δ=0.79, BF_+0_=6.42, *p*=.006) and trials with 20 dots (Δ=0.81, BF_+0_=3.13, *p*=.013). However, perceived numerosity in the two-group trials is on average *smaller* than in the one-group trials with intermediate dot size (Fig. 3, open circles in second graph). While the difference is not significant for trials with 16 dots (Δ=−0.43, BF_−0_=0.84, *p*=.064), there is extremely strong evidence for an effect in trials with 20 dots (Δ=−1.37, BF_−0_=12.1·10^2^, *p*<.001). These opposite effects can be coherently explained as an effect of dot size (“larger dots = smaller perceived numerosity”), but not as a pure grouping effect. If a grouping effect is present in these data – which we cannot rule out nor confirm – then it was obscured by a direct effect of dot size on perceived numerosity. In summary, while these results make it difficult to draw a conclusion about the effect of grouping by dot size on perceived numerosity, they do show clear evidence of feature-driven effects of dot size in stimuli with prolonged viewing, similar to previous findings using brief, static displays.

#### Spatial clustering

On average, subjects reported dots in two-group arrays as being more numerous than in one-group arrays (Fig. 3, left). Paired t-tests provide extremely strong evidence for this, both in trials with 16 dots (Δ=1.34, BF_+0_=33.2·10^2^, *p*<.001) and in trials with 20 dots (Δ=1.81, BF_+0_=35.1· 10^2^, *p*<.001). We see two possible explanations for this effect. First, it could be an effect of grouping: when dots are divided into two spatially separated groups, they are perceived as more numerous than when presented as a single group. However, this would be opposite to the grouping effects that we found in the conditions where dots were grouped by motion or color. An alternative explanation is that the effect is caused by a difference in the total size of the test array area between two-group trials (2×38=76 deg^2^) and one-group trials (64 deg^2^) (Fig. 2B, left). The current data do not allow us to distinguish between these two hypotheses – we will address this in Experiment 2.

#### Correlation analysis

We next examine whether there is evidence for correlations in effect sizes between the different features. To do so, we quantify the effect size for each subject as the trial-averaged percentage of overestimation in two-group arrays compared to one-group arrays (which we compute separately for each of the four grouping features). To illustrate this, let 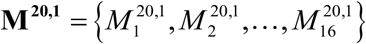 denote the responses in the 16 trials with a one-group motion stimulus and 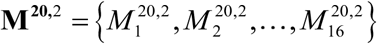 the responses in the 16 trials with a two-group motion stimulus. Then the average effect size is computed as

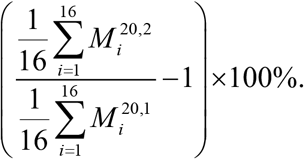

Hence, if a subject reported on average 19 dots on the two-group trials for this feature and 20 dots on one-group trials, then the effect size is 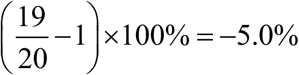 (i.e., two-group numerosities were 5% underestimated relative to one-group numerosities). For each subject, we computed the effect sizes separately for each of the four grouping feature and each of the two set sizes (16, 20). Thereafter, we averaged the effect sizes across the two set sizes, so that we end up with 4 effect size estimates per subject (1 per grouping feature).

The results of a Bayesian correlation analysis performed on these values (Table 1) reveal strong evidence for a correlation between color-based and motion-based grouping effects on perceived numerosity, but no other correlations. Results from a frequentist analysis are consistent with the Bayes Factors: the correlation between motion and color is significant (*p*=.004) and all others are non-significant (*p*>.21). This finding suggests that effects of color-based and motion-based grouping on numerosity estimates originate from a common, feature-invariant source, while the opposite effects found in conditions with grouping based on spatial clustering and dot size may have distinct origins.

**Table 1.** *Correlations between influences of the four grouping features on perceived numerosity*.

|  |  | **Spatial clustering** | **Size** | **Color** | **Motion** |
| --- | --- | --- | --- | --- | --- |
| Spatial clustering | Pearson's r | — |  |  |  |
|  | BF<sub>+</sub>0 | — |  |  |  |
| Size | Pearson's r | 0.146 | — |  |  |
|  | BF<sub>+</sub>0 | 0.498 | — |  |  |
| Color | Pearson's r | -0.172 | 0.111 | — |  |
|  | BF<sub>+</sub>0 | 0.079 | 0.364 | — |  |
| Motion | Pearson's r | 0.095 | 0.009 | 0.378 | — |
|  | BF<sub>+</sub>0 | 0.319 | 0.177 | 17.12 | — |

*Note .* For all tests, the alternative hypothesis specifies that the correlation is positive.

#### Interaction effects

While we are mainly interested in feature-driven and grouping-driven biases *within* features, for completeness we also perform a 2 × 2 × 4 (number of dots × number of groups × grouping feature) within-subjects ANOVA on the entire dataset (Table 2). The results indicate extremely strong evidence for an interaction between number of groups and grouping feature on perceived numerosity. Moreover, we find extreme evidence for main effects of both number of test dots and grouping feature, but no evidence for a main effect of number of groups. Finally, there is moderate evidence for an interaction between number of dots and grouping feature, with more dots resulting in larger differences.

**Table 2.** *Results from frequentist and Bayesian repeated-measures ANOVAs performed on the data from Experiment 1 (higher-order interactions are excluded)*.

| | Sum of Squares | df | F | p | BF<sub>inclusion</sub> | $\eta^2$ |
| --- | --- | --- | --- | --- | --- | --- |
| #Dots | 2582 | 1, 54 | 1077 | < .001 | $6.05 \cdot 10^{135}$ | .95 |
| #Groups | 13.8 | 1, 54 | 3.24 | .077 | 0.75 | .057 |
| Feature | 389 | 3, 162 | 38.0 | < .001<sup>a</sup> | $1.57 \cdot 10^{29}$ | .41 |
| #Dots × #Groups | 0.421 | 1, 54 | 0.42 | .521 | 0.13 | .008 |
| #Dots × Feature | 38.5 | 3, 162 | 6.72 | < .001 | 6.01 | .11 |
| #Groups × Feature | 211 | 3, 162 | 23.0 | < .001<sup>a</sup> | $2.20 \cdot 10^{13}$ | .30 |
| #Dots × #Groups × Feature | 8.30<sup>a</sup> | 3, 162 | 1.78 | .154<sup>a</sup> | 0.07 | .032 |

<sup>a</sup> Sphericity assumption was violated, but Greenhouse-Geisser and Huynh-Feldt corrections did not substantially change the results.

### Discussion

We find differences in perceived numerosity between two-group and one-group arrays in all four tested grouping features. However, there is variation across the tested features in both the effect direction and the conclusions that we are able to draw about the origin of the effects. Grouping by color and motion led in both cases to a decrease in the average perceived number of dots. For both features, we found evidence against feature-driven effects on perceived numerosity and no evidence in favor of such effects. Moreover, we found strong evidence for a correlation in the effect sizes between these two features. Altogether, these results suggest that the grouping effects in the color and motion conditions are caused by a shared, feature-invariant mechanism. By contrast, grouping by dot size and spatial clustering both *increased* perceived numerosity. However, for these two features we could not rule out that the effects may have been feature-driven rather than due to a difference in the number of groups. Indeed, in the dot-size conditions, we found strong evidence in favor of such effects. This does not exclude the possibility that grouping-driven effects were also present in those data, but we are at present unable to separate the two types of effect. In the spatial clustering conditions, the identified effect may have been caused by a difference in total array area between the two-group and one-group stimuli. Since Experiment 1 did not contain a manipulation of array area independent of the number of groups, we perform a second experiment to examine this possibility in more detail. A secondary aim of Experiment 2 is to test whether the evidence for the null effects for color and dot size are robust under small changes in experimental conditions.

## EXPERIMENT 2

### Methods

#### Availability of data and analysis files

The data and JASP analysis files related to this experiment are available at https://osf.io/vbdf5/.

#### Participants

Eighty-seven adults were recruited from the student population at the department of Psychology at Uppsala University (24.8±6.9 years of age, 59 females). Participants received either course credit or a cinema ticket for their participation

#### Stimuli and procedure

As in Experiment 1, subjects adjusted the number of dots in a response array to match the number of dots in a test array (Fig. 4A). The stimuli and procedure were the same as in Experiment 1, except for the following differences. The test arrays contained 10, 15, or 20 dots and each dot had a diameter of 0.20 deg. The visual properties of the test array were fixed throughout the experiment, while the total array area and dot size of the response array were varied in a 3 × 3 factorial design (Fig. 4B). Specifically, the total array area and dot size in the response arrays were half, the same, or double the area or size of the test array. Each subject was tested 3 times on each of the 27 combinations (3 numerosities × 3 array areas × 3 dot sizes). The order in which these 81 trials were presented was randomized per subject. For half of the subjects, the dots in the test array were yellow and the dots in the response array blue; for the other half, the reverse coloring was used. The color manipulation was included to verify that the null effect of color found in Experiment 1 can be replicated.

**Figure 4.**
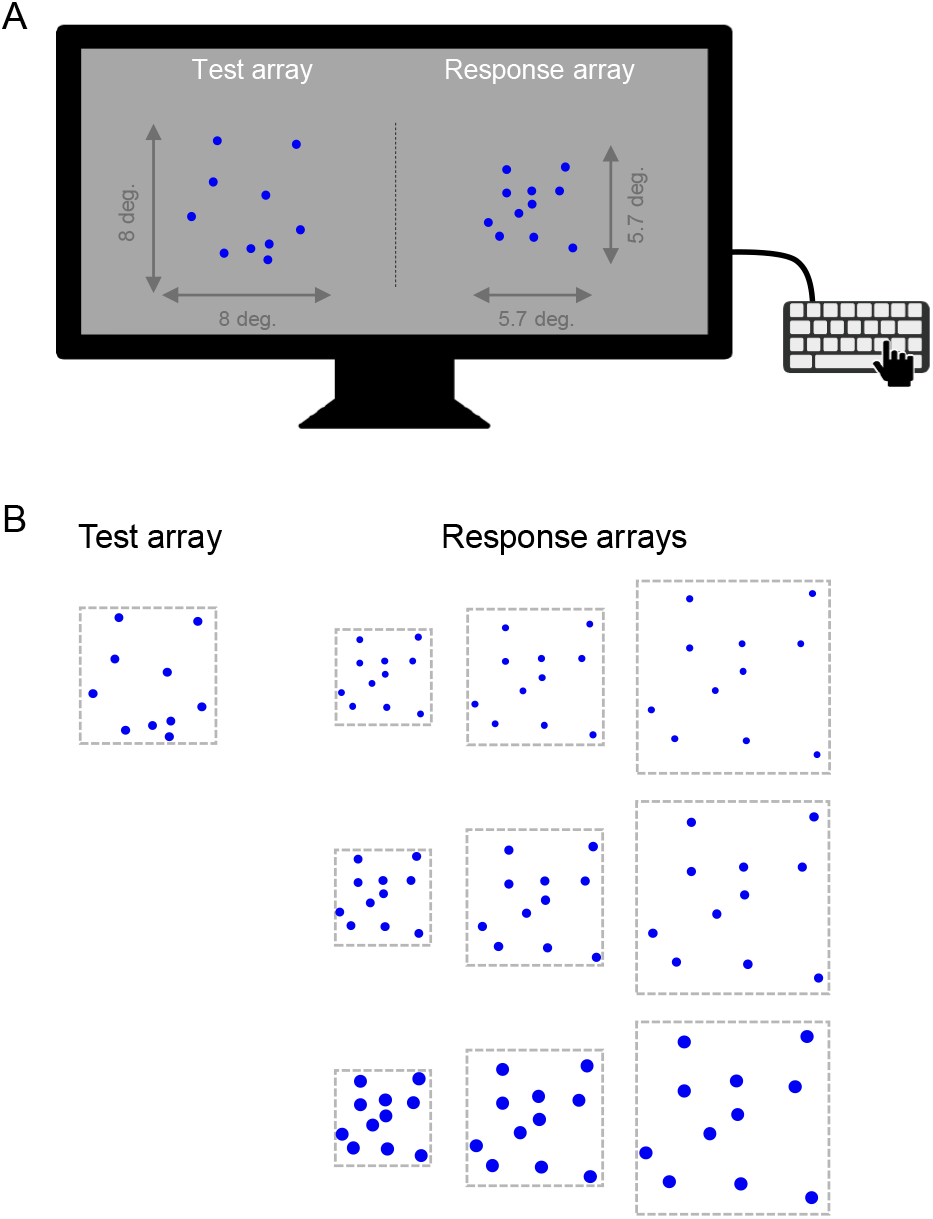
Illustration of stimuli used in Experiment 2. (A) A still image of an example stimulus in which the area of the response array is half that of the test array (32 *vs*. 64 deg^2^). (B) The dot size and area of the test array were fixed throughout the experiment, while the dot size and area of the response array were varied in a 3 × 3 factorial design.

### Results and discussion

We perform a 3 × 3 × 3 × 2 repeated-measures ANOVA with number of dots, array area, and dot size as within-subject factors, the color of the test dots as a between-subjects factor, and the reported number of dots as the dependent variable. The result reveals extremely strong evidence for a main effect of total array area on perceived numerosity (BF_inclusion_=∞, *p*<.001), with a positive relation: the larger the area of the response array, the smaller the setting of the number of response dots to obtain PSE with the test array (Fig. 5A). Hence, an increase in the response array area decreases the point of subjective equality with the test array, which means that the larger the area of an array, the larger the perceived number of dots. This is consistent with the results from Experiment 1, where spatial grouping of dots in the test array increased both the total array area and the average reported number of dots. Hence, it is plausible that the effect of spatial separation found in Experiment 1 was at least in part due to differences in array area, possibly on top of an effect due to a difference in the number of groups. Since we cannot disentangle these effects, it remains inconclusive what caused the increase in arrays with two spatial clusters compared to single-cluster arrays in Experiment 1.

**Figure 5.**
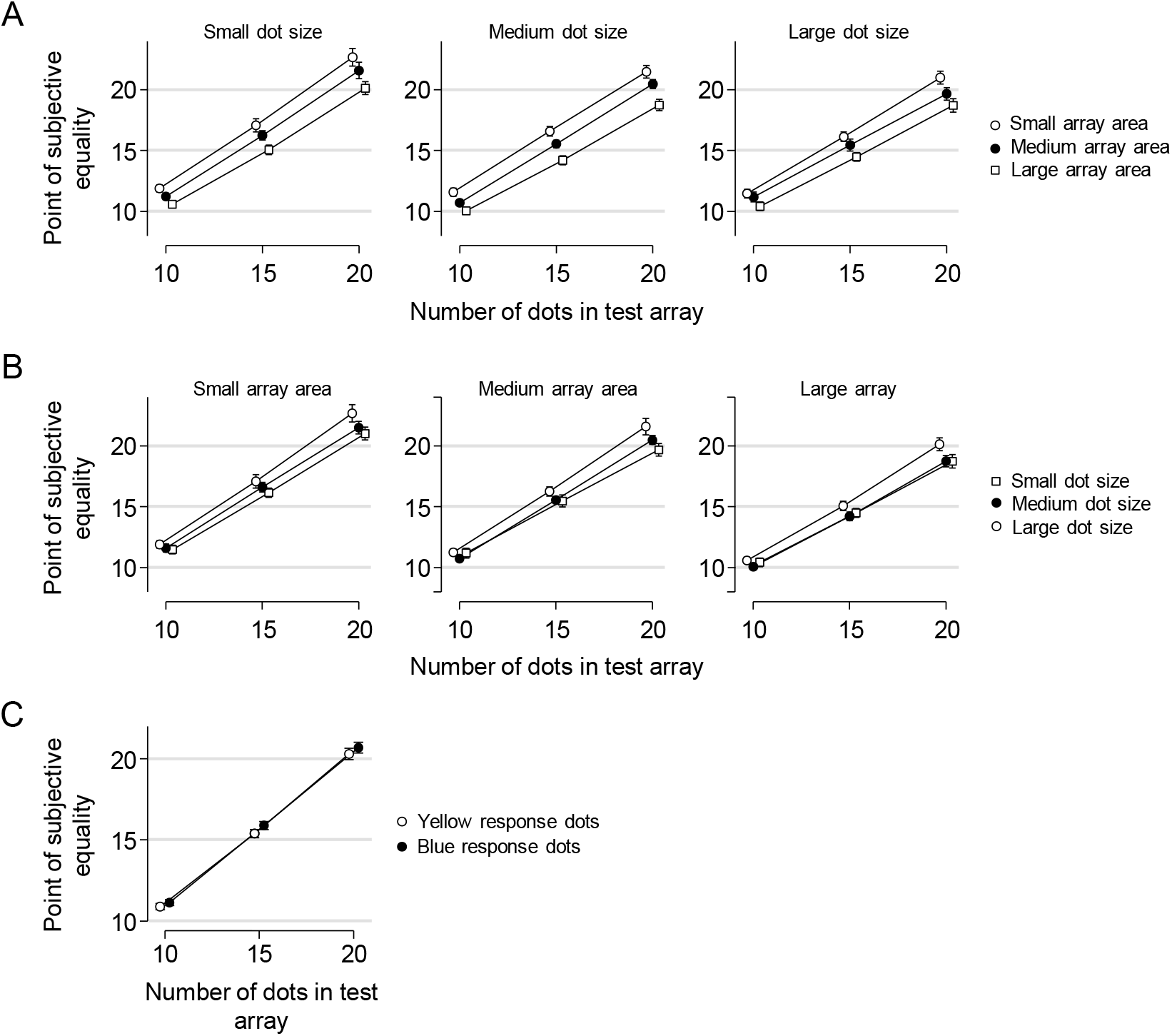
Results of Experiment 2. (A) Reported number of dots as a function of the number of dots in the test array, split by response array area (different lines) and response array dot size (different panels). Results are pooled across the two groups of subjects with different response dot colors. (B) An alternative visualization of the data that more clearly shows the effect of dot size (different lines). (C) The same data plotted separately for subjects with yellow and blue response dots (collapsed over dot size and array area conditions).

We also find extremely strong evidence for a main effect of the size of the dots in the response array on reported numerosity (BF_inclusion_=∞, *p*<.001), with a positive relation between the two variables: the smaller the dots in the response array, the smaller the reported number of dots (as indicated by the PSE, Fig. 5B). This result suggests that compensation occurs since the smaller dots in the response array are perceived as more numerous, which is consistent with the findings in Experiment 1 and provides further evidence that if there was a grouping-driven effect in the dot-size condition, then it was obscured by a feature-driven effect of dot size. Therefore, the results from the two-group dot-size condition in Experiment 1 are also inconclusive about the question how the number of groups affected perceived numerosity.

Furthermore, the ANOVA provides moderately strong evidence for the hypothesis that dot color does not affect the number of reported dots (Δ=0.38, BF_inclusion_=0.143; Fig. 5C), which is consistent with our finding in Experiment 1. This is further evidence that the color grouping effect was a purely grouping-driven effect. However, we should note that the corresponding *p* value is 0.019, which is conventionally interpreted as evidence *against* the null hypothesis.

Finally, it is worth mentioning that the results provide extremely strong evidence *against* an interaction effect between array area and dot size on perceived numerosity (BF_inclusion_=0.007, *p*=0.16), which suggests that these two biases act independently of each other. This finding is consistent with the lack of a correlation between effects of grouping by spatial clustering and grouping by dot size in Experiment 1.

## GENERAL DISCUSSION

### Summary

There has lately been much research on the human number sense and on the various biases in this cognitive ability. Most previous experimental work has used static stimuli with extremely brief exposure times. While that approach has delivered important insights into the human number sense, little is known about how the human number sense works in situations where stimuli are dynamic and viewed for extended periods. Here, we developed an experimental paradigm that more closely mimics such conditions and we used it to assess grouping effects in numerosity judgment. We found that grouping by color and motion decreased the average number of reported dots. Since we found evidence against direct influences of these features, we conclude that the effects were solely due to grouping. Moreover, we found a correlation between the effect sizes for these two features, which suggests that the effects may have a feature-invariant, higher-level origin. We were unable to draw conclusions about effects of grouping by spatial clustering or dot size, because these features were found to have a direct effect on perceived numerosity. Overall, our results suggest that perceived numerosity is negatively related to the number of groups, even though these effects may be overridden by stronger feature-driven effects in the opposite direction when grouping is established using spatial separate or object size as grouping feature. Therefore, the story told in Genesis, where Jacob spread a herd of sheep in smaller groups to appear more numerous [4], describes a hypothetical influence on perceived numerosity that is maybe not caused by grouping, but more likely due to the resulting larger total stimulus area formed by the spatially separate groups of sheep.

### Feature-driven effects on numerosity estimates

Our finding that smaller dots tend to be perceived as more numerous than larger dots is consistent with previous work that primarily used static stimuli with brief exposure [4,24–28]. It is currently unknown what the origin of this “smaller-dot-size-is-more” effect is. One possibility is that a given array area has more room for small items than for large items and that humans use this prior information during inference. However, other studies have found an opposite, “smaller-is-less” effect [20–23]. Findings from a recent eye-tracking experiment suggests that this effect may be related to attentional capture (Lindskog, Poom, & Winman, in preparation).

Our results also revealed an effect of total array area on perceived numerosity: the larger the area, the larger the average perceived numerosity. This result is also in line with previous studies [7,14,15]. A speculative explanation of the area-based bias is that when humans make numerosity judgments, they take into account their previous experiences. If a person has experienced that large areas or volumes tend to contain more objects than small ones and that smaller objects are typically more abundant within a given area or volume, then this could create a bias similar to the one we found. Consistent with this speculation, is has been found that increasing the size of a container increases the perceived number of beans in the container [35]. Moreover, this type of explanation is consistent with findings that biases may be reduced through practice [27] and instructions given to the observers [36]. Hence, the effect of area size on numerosity estimates may be partly due to top-down processes. However, there also is evidence for low-level, bottom-up influences of area on perceived numerosity. In particular, a recent study found that adaptation to a size stimulus alters subjects’ numerosity estimates: adapting to a larger size reduces the perceived numerosity in a subsequent stimulus (and vice versa) [37], which is consistent with our finding that larger areas are perceived as more numerous. It has to be kept in mind, however, that stimuli were viewed for prolonged durations in our experiment. Since adaptation effects are typically transient, it remains to be seen whether results from adaptation studies generalize to our kind of stimulus.

Finally, theories about visual crowding [38,39] may offer a single explanation of both the effect of dot size (“smaller dots is more”) and the effect of array area (“larger area is more”) that we found in our data. When objects are very close to each other, they are difficult to distinguish – especially in the periphery – which could lead to a reduction in perceived numerosity. Hence, the increase in perceived numerosity when making the dots smaller or the area larger might be due to improved distinguishability.

### Limited-dot-lifetime displays

Besides contributing further empirical insights into the number sense, the present study also introduced a novel experimental paradigm, in which stimuli were dynamic and viewed for prolonged durations. Our main motivation for using this kind of stimulus was that we believe that such stimuli are representative for a large class of numerosity judgments outside the laboratory. However, another advantage of LDDs is that they provide a more accurate percept of the total array area, thus providing the experimenter with stronger control over this experimental variable. The experiments presented here do not address the issue whether numerosity judgment tasks with LDDs target the same brain mechanisms as the traditional paradigm with brief and static stimuli. Therefore, it could be informative if future studies would directly compare empirical properties of number estimation in brief, static stimuli on the one hand and dynamic stimuli with prolonged viewing on the other hand. For example, some studies have argued that numerosity estimates are mainly derived from low-level visual cues [40,41]. It is unclear whether the use of longer presentations would increases or decreases the tendency to use visual cues. On the one hand, longer presentation times are expected to reduce the influence of rapid, bottom-up effects caused by basic features. However, on the other hand, they may also lead to more accurate estimates of the visual features, which could be a reason to rely on them more strongly. Since our paradigm allows for the use of arbitrary stimulus times, without the risk that subjects will explicitly count the stimuli, future studies could use tasks with LDDs obtain more insight into the relation between stimulus duration and the use of visual cues in number judgment. Moreover, it could be interesting to test whether the putative link between numerosity judgment accuracy and mathematical ability [1] generalizes to tasks with LDDs. Finally, other psychophysical methods than the adjustment procedure can be used with LDDs, such as the 2AFC method using fixed presentation times. Such procedures using LDD could also be useful when prolonged viewing is required, for example in functional imaging studies.

### Interactions between density and numerosity

A limitation of the present study is that numerosity in Experiment 1 was near-perfectly correlated with density: the larger the number of dots, the higher dot density. Therefore, we cannot rule out that subjects in this experiment judged density rather than numerosity. The results from Experiment 2 may provide some information on this question, because there we varied array area (and, thus, density) independently of numerosity. Our finding that larger array area prompted larger numerosity estimates is opposite to what we would expect if density had been used as a cue for numerosity. Therefore, it seems that, at least in Experiment 2, subjects judged numerosity largely independently of density. This is consistent with previous studies claiming that the two types of judgment are based on separate mechanisms (e.g., [37,42,43]). However, it should be mentioned that there is also evidence suggesting the opposite, namely that human sense of numerosity and density are intimately intertwined (e.g., [44,45]). As far as we are aware, all previous work on the relation between numerosity and density judgments is based on experiments with static stimuli that were viewed for extremely short times. It would be worthwhile to assess how those results translate to a context with dynamic stimuli that can be viewed for prolonged periods.

## ACKNOWLEDGEMENTS

LP acknowledges support from the Swedish Research Council (Vetenskapsrådet; reg. nr. 2013-01005). RB acknowledges support from the Swedish Research Council (Vetenskapsrådet; reg.nr. 2015-00371) and Marie Sklodowska Curie Actions, Cofund (project INCA 600398).

## Footnotes

a Subjects rarely reported numbers close to the edges of the range; over 98% of the responses were in the range [10, 30].

b We do not include higher-order interactions in these analyses (by selecting the option “Across matched models” in JASP), because this provides a better comparison with p-values of main effects from a frequentist ANOVA.

